# Chromosome-scale genomes of *Dugesia* reveal contrasting modes of genome size evolution and structural stability in flatworms

**DOI:** 10.64898/2025.12.11.693146

**Authors:** Daniel Dols-Serrate, Ignacio Tenaguillo-Arriola, Vadim A. Pisarenco, Marta Olivé-Muñiz, Julio Rozas, Marta Riutort

**Affiliations:** Departament de Genètica, Microbiologia i Estadística, Facultat de Biologia, Universitat de Barcelona, Av. Diagonal 643, 08028, Barcelona, Catalonia, Spain; Institut de Recerca de la Biodiversitat (IRBio), Universitat de Barcelona, Av. Diagonal 643, 08028, Barcelona, Catalonia, Spain

**Author notes:** Corresponding authors: M. Riutort, Daniel Dols-Serrate and Julio Rozas.

**Keywords:** *Dugesia* flatworms, PacBio, genome assemblies, chromosomal rearrangements, genome size evolution, *Schmidtea mediterranea*

## Abstract

High-quality, chromosome-scale genomes are crucial for understanding biological processes, yet many metazoan lineages, including most Lophotrochozoa, remain underrepresented in genome databases. Among these, planarians (Platyhelminthes, Tricladida), particularly *Dugesia*, are a globally distributed and phenotypically diverse group that has become an important model in evolutionary biology, notably for investigating the genetic effects of agametic asexuality. However, the lack of chromosome-scale assemblies has limited progress. Here, we present the first chromosome-scale genomes of four Western Mediterranean *Dugesia* species, displaying the first intra- and intergeneric comparisons. Comparison with the regeneration model organism *Schmidtea mediterranea*, rejects a whole-genome duplication as the cause of differences in chromosomal number and genome size between genera. Instead, Dugesia shows extensive lineage-specific and differential expansions of DNA transposable elements, likely contributing to genome size variation during diversification. Despite differences in the dynamics of structural genome rearrangements observed between genera, both groups lack the conservation of ancestral metazoan linkage groups, supporting the idea that genome structural instability is a key feature of flatworm genome evolution. Our newly generated genomic resources and findings offer vital insights into the genetic basis of diversification and establish *Dugesia* as a valuable model for studying metazoan genome dynamics, including the evolution of alternative reproductive systems.

## 1. Introduction

Biological evolution proceeds through cumulative genomic changes that shape genetic diversity and weave the continuous and complex process that ultimately drives speciation. The advent of next-generation sequencing (NGS) has greatly advanced comparative genomics, deepening our understanding of how genomes evolve, regulate biological functions, and give rise to phenotype diversity^1,2,3,4,5^. Chromosome-scale genome assemblies, in particular, are key for elucidating the mechanisms and evolutionary history of structural variation, chromosomal rearrangements (CRs), as well as their consequences for reproductive isolation and speciation^6,7,8,9,10,11^. Although such high-quality genome resources are becoming increasingly available for non-model organisms, their distribution across the Tree of Life remains uneven, with significant taxonomic biases^12,13,14^. This disparity is particularly striking in Lophotrochozoa^15^, a major animal superphylum that is severely underrepresented in genomic databases. Despite accounting for approximately 40% of all metazoan phyla^16,17^—including major groups such as Mollusca, the second-largest metazoan phylum—the number of available Lophotrochozoa genomes stands in stark contrast to their immense biodiversity, with less than 0.01% of known species sequenced to date^18,19^. This gap poses a challenge to the field of evolutionary biology and limits our ability to identify innovations unique to this vast portion of animal life, which in turn hinders our comprehensive understanding of metazoan biology. Recent genomic surveys have already underscored the importance of filling this gap, revealing unexpected insights into lophotrochozoan genome architecture and plasticity^20,21^. Planarians (Platyhelminthes, Tricladida), the focus of this study, exemplify this taxonomic imbalance and remain among the most historically overlooked groups in terms of genomic resources.

Planarians are a species-rich taxon encompassing thousands of species distributed worldwide and exhibiting a rich phenotypic diversity across terrestrial, marine, and freshwater environments^22^. Flatworms have been extensively studied in ecology^23^, autophagy dynamics^24^, and biogeography^25,26^, to name a few disciplines. However, planarians are best known for their unmatched regenerative abilities^27,28,29^, enabled by their pluripotent stem cells (neoblasts), which have established them as key model systems in regeneration and stem-cell biology. It would be reasonable to expect a wealth of genomic resources for planarians, particularly given that the model species *Schmidtea mediterranea* Benazzi, Baguñà, Ballester, Puccinelli & Del Papa, 1975, was among the early cohort of genomes sequenced using the Sanger-sequence technology^30,31^. Despite this, only five triclad genomes are currently available: those of the four *Schmidtea* species^27^, and that of *Dugesia japonica* Ichikawa & Kawakatsu, 1964^32^. Notably, *Schmidtea* genomes have revealed unexpected and peculiar features, including extensive genome scrambling and a complete absence of conserved ancestral metazoan linkage groups (MALGs) across species, prompting the proposal that genome-wide structural instability may be a hallmark of flatworm genome evolution^27^.

The genus *Dugesia* Girard, 1850, closely related to *Schmidtea* (Figure 1), comprises approximately 110 freshwater species, roughly half of the family Dugesiidae Ball, 1974, and exhibiting a broad global distribution. In recent years, *Dugesia* has emerged as a model system in evolutionary biology, owing to the interplay between its exceptional species richness in the geologically dynamic Mediterranean region, and the presence of distinct biotypes either different reproductive modes (sexual or asexual by fission) and/ or ploidy levels within species (i.e., sexual diploids, fissiparous triploids, and multiple polyploidies or aneuploidies). Extensive research in the Western Mediterranean has leveraged these traits, leading to the discovery of numerous new species, the identification of a unique putative hybridization-driven speciation event, frequent chromosomal translocations in asexual populations, evidence for the mosaic-Meselson effect in fissiparous individuals, and the identification of long-lived asexual lineages^33,34,35,36,37^. These efforts have also established a well-resolved phylogeny linking diversification patterns to abiotic events, progressively advancing from the use of a few molecular markers to reduced-representation genomic datasets, and transcriptomes. Despite its importance and this extensive body of work, the only genomic reference available for *Dugesia* is a non-chromosome-level draft-assembly of *D. japonica*, comprising 10,017 scaffolds^32^.

**Figure 1.**
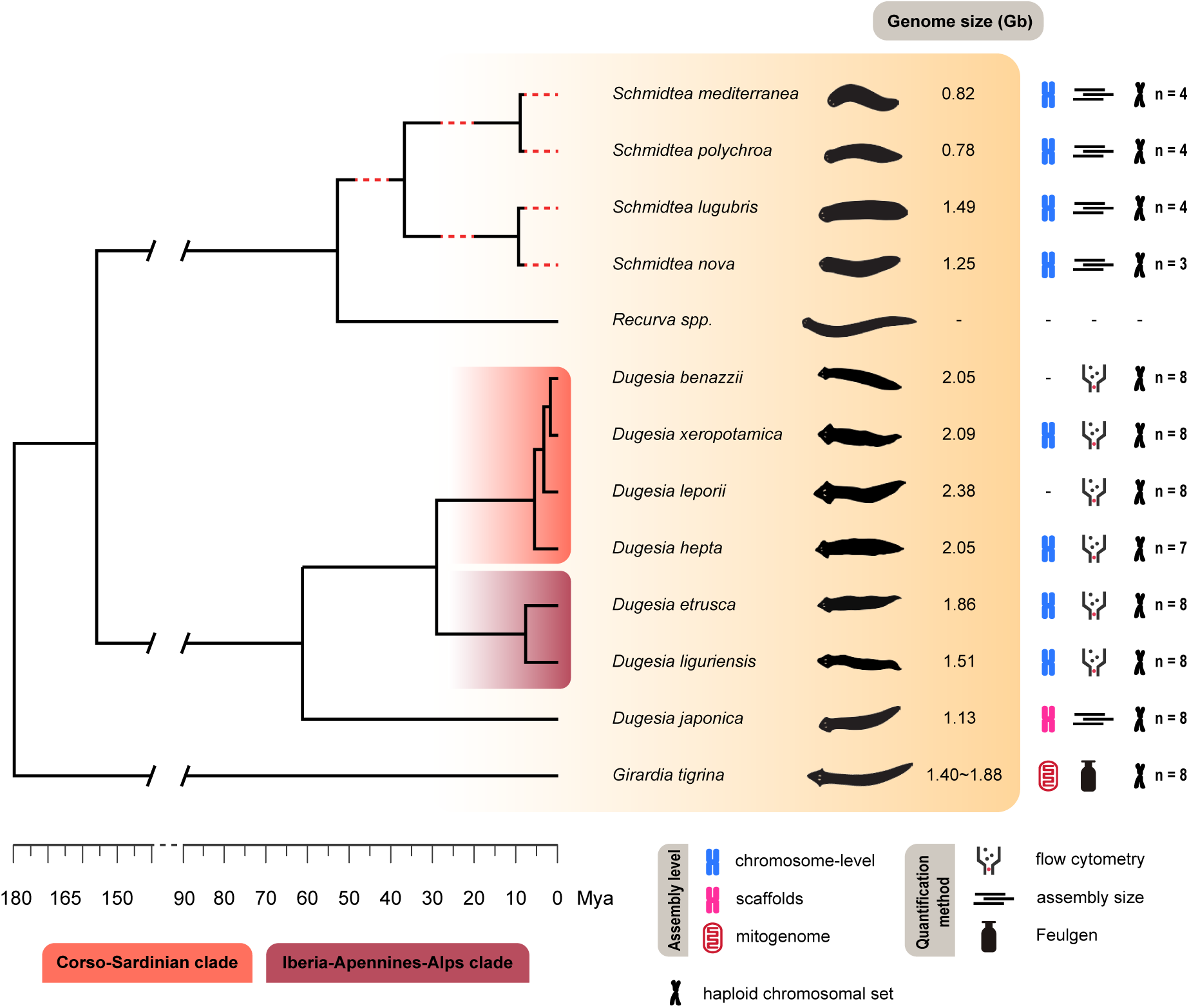
Schematic time-calibrated tree illustrating the phylogenetic relationships among species of *Dugesia*, *Girardia*, *Recurva*, and *Schmidtea* based on Leria et al.^108^ and Solà et al.^26^. Red dotted lines indicate branches for which the time to their most recent common ancestor (MRCA) is unknown.

Generating high-quality genomes for planarians is exceptionally challenging due to three key factors: i) the limited number of already sequenced genomes with the resulting poor understanding of their genomic architecture; ii) the large genome sizes coupled with a high abundance of TEs, which complicate assembly; and iii) the pronounced A/T nucleotide bias, which further hinders sequencing and assembly. Consequently, it is not surprising that, despite their significance in evolutionary biology, planarians remain vastly unrepresented in genomic databases. This critical gap hinders deeper investigation into the molecular mechanisms underlying diversification, as well as the evolutionary significance and long-term fate of sexual and asexual lineages, and more generally limits the potential of planarians as model organisms in Lophotrochozoa evolution.

To address this gap, we generated chromosome-scale genomes of four sexual diploid *Dugesia* species (Figure 1): two endemic insular species from the Corso-Sardinian clade, viz. *Dugesia hepta* Pala, Casu & Vacca, 1981, and *Dugesia xeropotamica* Stocchino, Dols-Serrate & Riutort, 2025, and two continental ones from the Iberia-Apennines-Alps clade, viz. *Dugesia etrusca* Benazzi, 1946, and *Dugesia liguriensis* de Vries, 1988.

Additionally, we also report the genome size (GS) estimates for two other Sardinian endemics for which genome assembling was not feasible, viz. *Dugesia benazzii* Lepori, 1951, and *Dugesia leporii* Pala, Stocchino, Corso, & Casu, 2000.

These new genomic resources enable us to: (i) investigate the genomic evolution within the genus, including the role of chromosomal rearrangements (CRs) in speciation (Figure 1); (ii) to explore whether a whole-genome duplication (WGD) could underlie the differences in chromosome number and GS between *Schmidtea* and *Dugesia*, considering both the higher GS and the doubled haploid complement in Western Palearctic *Dugesia* species (Figure 1); (iii) assess whether genomic patterns recently described in *Schmidtea,* including the extensive genome-wide rearrangements and the complete lack of conservation of MALGs^27^, are shared by *Dugesia* species. By generating these genomic resources and addressing the three aims outlined here, our study substantially advances population and evolutionary genomics in these two model groups and provides new perspectives on metazoan genome evolution.

## Results

### 2.1. New *Dugesia* genomes: assembly and annotations

We generated chromosome-level genome assemblies for four selected *Dugesia* species. Since wild populations of the selected continental species are known to present coexisting biotypes, we characterised them via karyological analysis and flow cytometry to ensure the selection of only diploid individuals. Subsequently, we generated the draft genomes for our species, referred below as Detr_v1 (*D. etrusca*), Dlig_v1 (*D. liguriensis*), Dhep_v1 (*D. hepta*), and Dxer_v1 (*D. xeropotamica*), using Pacific Biosciences’ CLR (Dhep_v1 & Detr_v1) and HiFi (Dlig_v1 & Dxer_v1) reads and Hi-C for scaffolding. The haploid genome assembly sizes yielded between 2.09 and 2.96 Gb (Table 1). Overall, N50 and L50 values, alongside Hi-C contact maps, indicate high contiguity as well (Table 1; Supplementary Fig. S1; Supplementary Fig. S2). Contact maps showed that the number of largest scaffolds per assembly (two or more orders of magnitude larger than any other given scaffold) was equal to the known haploid chromosomal number of each species^38,39,40^. These scaffolds, hereafter referred to as pseudochromosomes, represent between 52.5% and 82.7% of their respective total assembly GSs (Table 1; Supplementary Fig. S2), which in three species yield bigger estimates than those from flow cytometry. Genome completeness was assessed using BUSCO^41^, yielding results comparable to those reported for other freshwater planarians (Table 1; Supplementary Fig. S3; cf. Ivanković et al.^27^). This strengthens the idea that BUSCO may be sub-optimal for planarian genomes owing to their substantial loss of genes^42^, highlighting the need for a group-specific BUSCO set (e.g., ^27,32,36^), as reflected in the high number of missing genes (Supplementary Table S1). However, BUSCO scores improved after using the structurally annotated assemblies (Table 2; Supplementary Fig. S3).

**Table 1.** Genome assembly statistics after contaminants removal.

|  | **Detr_v1** | **Dhep_v1** | **Dlig_v1** | **Dxer_v1** |
| --- | --- | --- | --- | --- |
| Flow cytometry (Gb) | 1.86 | 2.05 | 1.51 | 2.09 |
| Assembly size (bp) | 2,961,733,428 | 2,090,052,221 | 2,136,556,669 | 2,484,242,095 |
| GC content (%) | 27.22 | 27.64 | 27.34 | 27.94 |
| Number of scaffolds | 59,495 | 15,220 | 8,935 | 35,214 |
| Longest scaffold (Mb) | 288.8 | 335.3 | 299.4 | 321.6 |
| Sequence in chromosomes (%) | 52.50 | 82.73 | 74.26 | 69.18 |
| N50 (Mb) | 141.2 | 242.1 | 162.2 | 176.5 |
| L50 | 8 | 4 | 5 | 6 |
| Complete BUSCOs |  |  |  |  |
| Metazoa data set | 553 (58.0%) | 616 (64.6%) | 659 (69.1%) | 556 (58.3%) |
| Arthropoda data set | 599 (59.1%) | 591 (58.3%) | 636 (62.7%) | 597 (58.9%) |
| Eukaryota data set | 189 (74.1%) | 180 (70.6%) | 199 (78.0%) | 188 (73.7%) |

**Table 2.** Genome annotation statistics.

|  | **Detr_v1** | **Dhep_v1** | **Dlig_v1** | **Dxer_v1** |
| --- | --- | --- | --- | --- |
| Repetitive regions (%) | 78.32 | 78.68 | 78.91 | 83.29 |
| Transposable & repetitive elements (bp) | 2,318,718,339 | 1,643,800,512 | 1,685,580,944 | 2,067,343,575 |
| Number of elements | 5,922,150 | 4,067,511 | 4,794,532 | 5,109,694 |
| Genome annotation |  |  |  |  |
| Protein-coding genes | 53,829 | 35,435 | 45,367 | 39,727 |
| Number of transcripts | 56,813 | 37,773 | 47,772 | 41,966 |
| Supported genes | 27,240 | 18,588 | 21,268 | 29,282 |
| Functionally annotated genes | 39,543 | 28,146 | 34,991 | 30,520 |
| Number of tRNA | 1,336 | 6,106 | 19,549 | 11,762 |
| Number of tRNA after filtering | 842 | 3,247 | 16,080 | 4,258 |
| Complete BUSCOs |  |  |  |  |
| Metazoa data set | 699 (73.3%) | 689 (72.2%) | 539 (56.5%) | 732 (76.7%) |
| Arthropoda data set | 718 (70.9%) | 701 (69.2%) | 561 (55.3%) | 736 (72.7%) |
| Eukaryota data set | 211 (82.8%) | 205 (80.4%) | 170 (66.7%) | 216 (84.7%) |

Structural annotation using BRAKER2 revealed 35,345-53,829 protein-coding genes in *Dugesia* (Table 2), far exceeding the number reported in *D. japonica* (12,031 genes; cf. Tian et al.^32^). Gene counts for *D. hepta* and *D. xeropotamica* were instead similar to those reported in *S. mediterranea* (∼31,000 genes in Grohme et al.^42^; 34,976 genes modelled using Guo et al.^43^ assembly). Sequence similarity-based searches indicated that most protein-coding genes (73.5-79.4%; Table 2) had identified matches in the surveyed protein databases.

### 2.2. Genome size pattern differences between *Schmidtea* and *Dugesia*

#### 2.2.1. Whole-genome duplication

We hypothesized that a WGD event might be the underlying cause for the notable karyotypic differences between both genera and also the GS disparity between *S. mediterranea* and *Dugesia*. We explored this scenario using three approaches: (1) modelling the *K*_s_ distribution of each *Dugesia* assembly’s paranome using wgd v2^44^, (2) assessing the intragenomic collinearity in the assemblies using MCScanX^45^, and (3) analysing macrosynteny patterns. All lines of evidence reject a WGD: *K*_s_ distributions show an exponential decay consistent with mostly recent duplications (Figure 2a; Supplementary Fig. S4), collinearity is very low (1.40% in Detr_v1, 1.04% in Dhep_v1, 2.19% in Dlig_v1, and 0.40% in Dxer_v1), and macrosynteny (Figure 3b) shows no *Schmidtea* syntenic blocks matching multiple *Dugesia* blocks.

**Figure 2.**
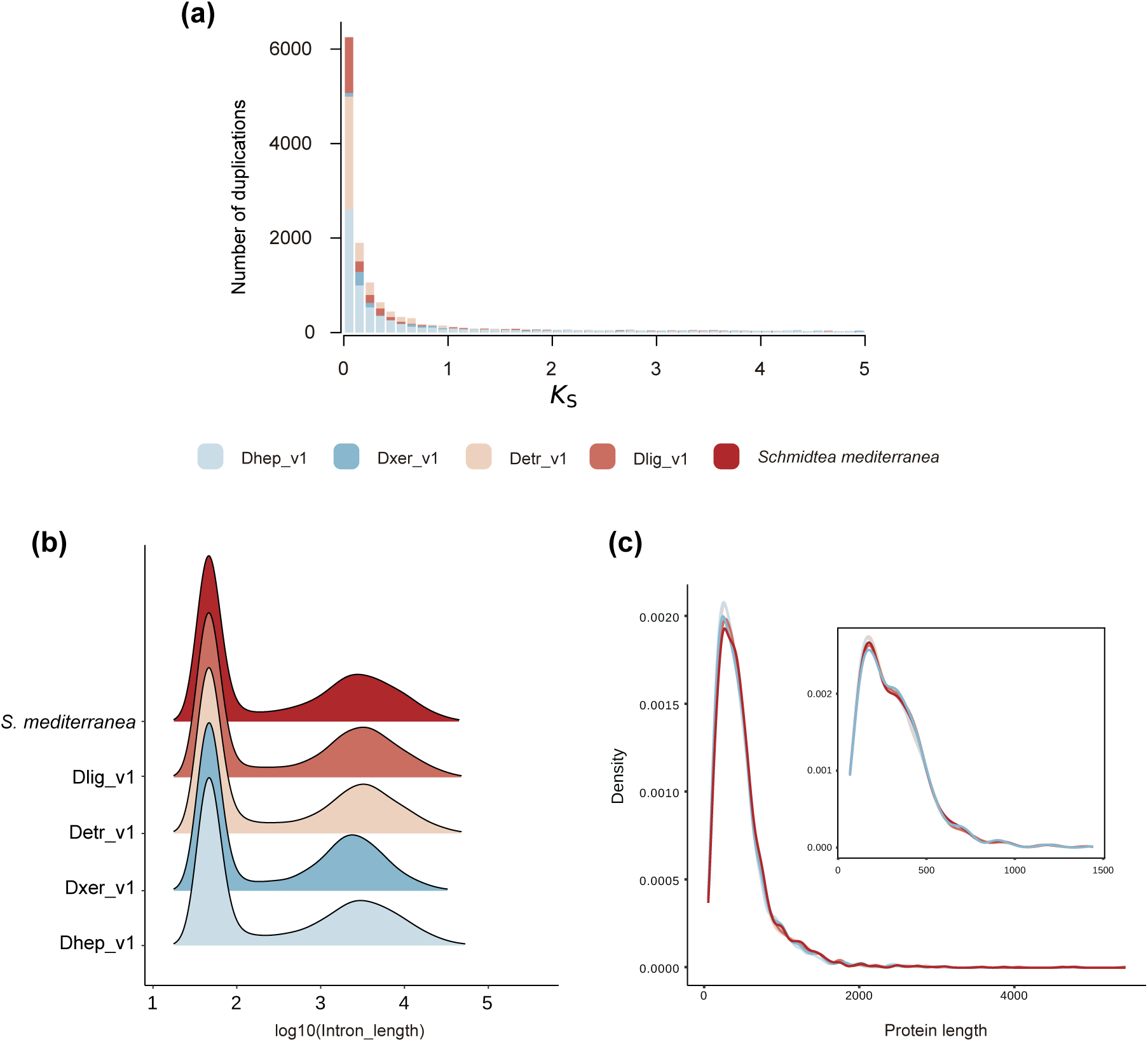
(a) Overlayed paranome *K_s_* distributions of the four *Dugesia* genome assemblies showing an exponential decay pattern. **(b)** Ridgeline plots of intron length distributions for all introns comprised in the 2,266 single-copy orthogroups shared among *D. hepta* (Dhep_v1), *D. etrusca* (Detr_v1), *D. liguriensis* (Dlig_v1), *D. xeropotamica* (Dxer_v1), and *S. mediterranea*. **(c)** Density distribution of CDS lengths of all 2,266 single-copy orthologues. The inset shows the distribution for the 571 exon-equivalent orthologues.

**Figure 3.**
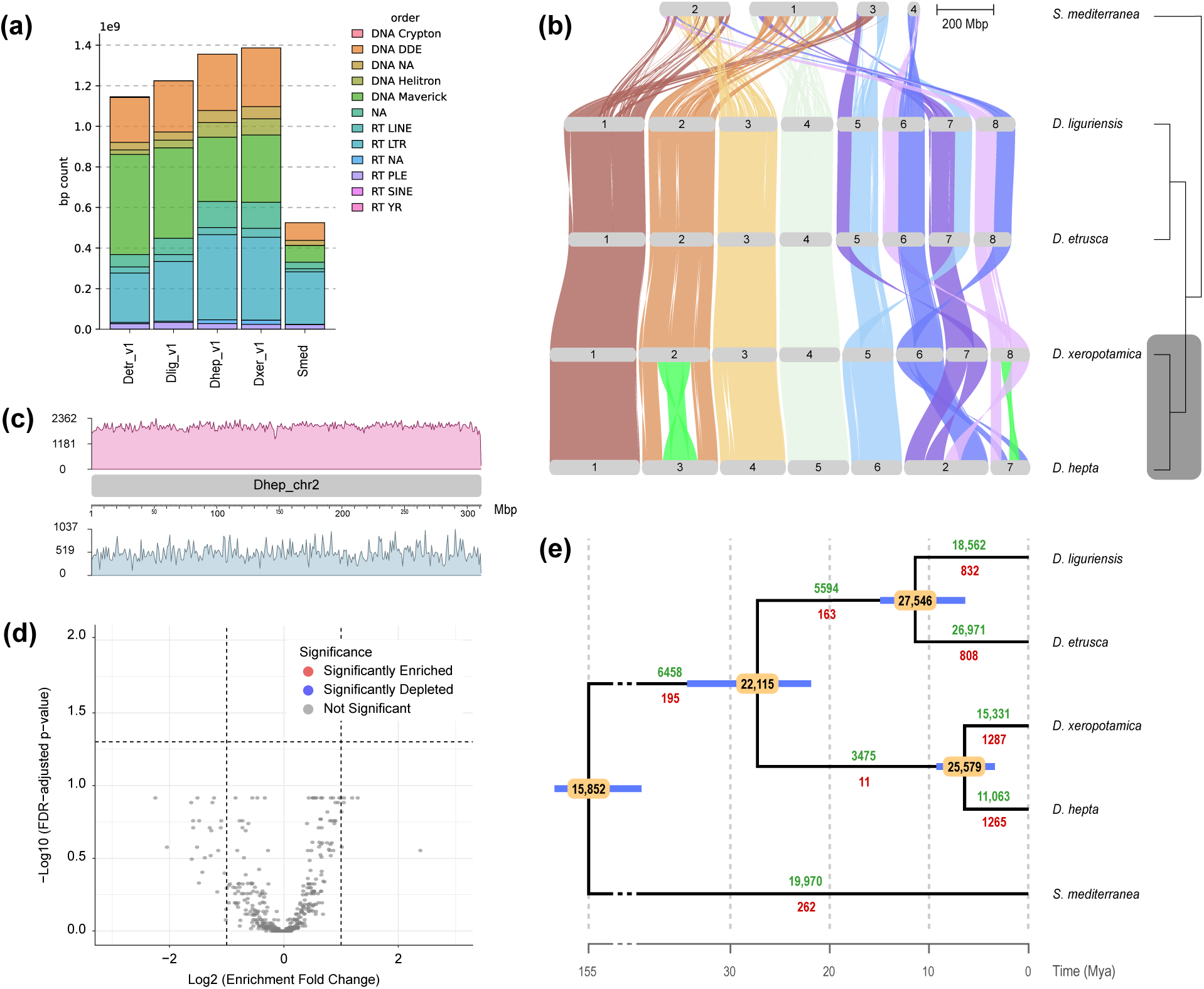
(a) Stacked histograms showing the total repeat content (bp) of each genome assembly, classified by TEs orders. **(b)** Synteny analysis between the four *Dugesia* species and *S. mediterranea*, with ribbons colored based on *D. xeropotamica* (n = 8). Pericentric inversions on chromosomes 2 and 7 are highlighted in green. Right: cladogram depicting phylogenetic relationships. Highlight in dark grey identifies the Corso-Sardinian clade. **(c)** Density plot of TEs (pink) and genes (cyan) along chromosome 2 of *D. hepta*. **(d)** Vulcan plot showing the enrichment analysis for 20 kb windows flanking synteny breakpoints across *D. hepta*’s chromosome 2. Horizontal dotted line indicates the statistical significance threshold. Vertical dotted lines represent a 2-fold depletion (left) and a 2-fold enrichment (right). **(e)** Time-calibrated phylogeny of the studied species showing the minimum number of gene gains (green) and losses (red) estimated with BadiRate. Ancestral gene counts are shown within yellow bubbles on top of the tree nodes, and blue bars represent the maximum and minimum time estimates for each node.

#### 2.2.2. Changes in intron and CDS lengths

To determine the genomic features underlying the GS differences between *Dugesia* and *S. mediterranea*, we analysed intron and CDS length distributions (Figure 2b-c; Table 3) of single-copy orthologues (SOs). These SOs were identified with Broccoli v.1.2^46^, yielding 2,266 shared genes between *Dugesia* and *S. mediterranea*, 571 of them being exon-equivalent (exhibiting the same number of exons in all species).

**Table 3.** Summary length (bp) statistics of introns and protein-coding sequences (CDS) of single-copy orthologues.

| **All introns** | **Detr_v1** | **Dhep_v1** | **Dlig_v1** | **Dxer_v1** | **Smed** |
| --- | --- | --- | --- | --- | --- |
| Number of introns | 11,032 | 10,692 | 10,370 | 10,312 | 9,273 |
| Shortest intron | 33 | 38 | 38 | 33 | 34 |
| Median lenght | 94 | 86 | 123 | 107 | 96 |
| Mean length | 3326 | 3398 | 3375 | 3253 | 2435 |
| Longest intron | 133,684 | 114,161 | 90,611 | 132,065 | 248,596 |
| **Short introns ($\leq 316$ bp)** |  |  |  |  |  |
| Number of introns | 6060 | 5879 | 5527 | 5519 | 4999 |
| Shortest intron | 33 | 38 | 38 | 33 | 34 |
| Median lenght | 47 | 47 | 47 | 46 | 47 |
| Mean length | 59.75 | 58.72 | 59.02 | 58.08 | 58.17 |
| Longest intron | 316 | 316 | 315 | 316 | 316 |
| **Large introns ($> 316$ bp)** |  |  |  |  |  |
| Number of introns | 4972 | 4813 | 4843 | 4793 | 5671 |
| Shortest intron | 318 | 317 | 320 | 317 | 317 |
| Median lenght | 4165 | 4041 | 4140 | 3866 | 3118 |
| Mean length | 7307 | 7476 | 7159 | 6932 | 5215 |
| Longest intron | 133,684 | 114,161 | 90,611 | 132,065 | 248,596 |
| **All CDS** |  |  |  |  |  |
| Number of CDS | 2266 | 2266 | 2266 | 2266 | 2266 |
| Shortest CDS | 52 | 67 | 67 | 62 | 67 |
| Median length | 342 | 340.5 | 355 | 350 | 364 |
| Mean length | 423.6 | 417.1 | 437.5 | 433.3 | 445.2 |
| Longest CDS | 4196 | 4052 | 4807 | 4252 | 4224 |
| **Exon-equivalent CDS** |  |  |  |  |  |
| Number of CDS | 571 | 571 | 571 | 571 | 571 |
| Shortest CDS | 71 | 71 | 71 | 71 | 67 |
| Median length | 266 | 267 | 273 | 275 | 276 |
| Mean length | 299.4 | 300.3 | 304.5 | 305 | 306.4 |
| Longest CDS | 1336 | 1349 | 1351 | 1349 | 1379 |

Based on intron length distributions (Figure 2b; Supplementary Fig. S5), we set a threshold value of 2.5 for the logarithmic intron length (316 bp), and classified introns as small-sized (≤ 316 bp) or large-sized (> 316 bp). Small introns showed little difference among *Dugesia* species and *S. mediterranea*, although the latter had lower numbers. Large introns, in contrast, were longer although less abundant in *Dugesia* (Table 3; Figure 2b). In contrast, CDS length distributions were largely similar across species (Figure 2c; Table 3). Overall, since introns represent a small fraction of the genome, our results indicate that GS variation across the genus primarily involves, as expected, changes in intergenic regions.

### 2.3. Macrosynteny and CRs

We inferred and visualized macrosynteny blocks using GENESPACE v.1.2.3^47^. Overall, macrosynteny is well preserved among the four *Dugesia* species (Figure 3b), although rearrangements were detected on chromosomes 5 to 8 between the Corso-Sardinian and the Iberia-Apennines-Alps clades. Within the latter clade, no major translocations or inversions were observed (Figure 3b) despite that the species shared their last common ancestor ∼9.5 Mya^26^. As expected, *D. hepta* exhibits major translocations and fusions, affecting chromosomes 2 and 7, with chromosome 2 sharing major syntenic blocks with multiple chromosomes in the other species (*D. xeropotamica*’s chromosomes 6, 7, 8, and chromosomes 5, 6, 7, and 8 of *D. etrusca* and *D. liguriensis*) (Figure 3b). These findings contrast with observations in *Schmidtea*, where loss of synteny is pervasive among congeners^27^. Finally, ODP analyses confirm that, as in *Schmidtea*, *Dugesia* shows no detectable synteny conservation of MALGs as well (Figure 4).

**Figure 4.**
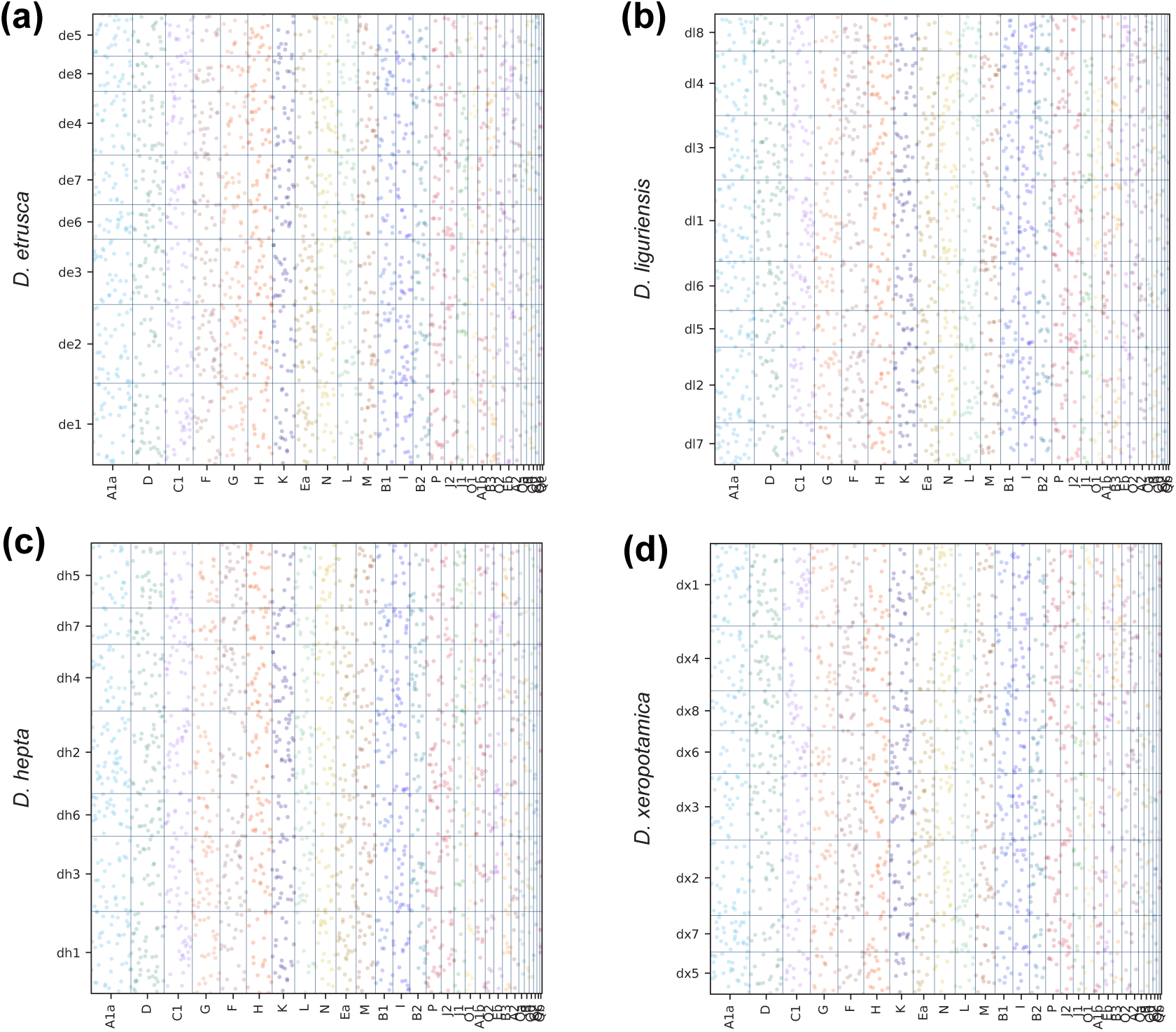
Dotplots showing the relationship between metazoan ancestral linkage groups (MALGs) and **(a)** *D. etrusca*, **(b)** *D. liguriensis*, **(c)** *D. hepta*, and **(d)** *D. xeropotamica*. MALGs without significant enrichment are plotted in light colors. No MALGs enrichment was detected in any *Dugesia* species.

**Figure 5.**
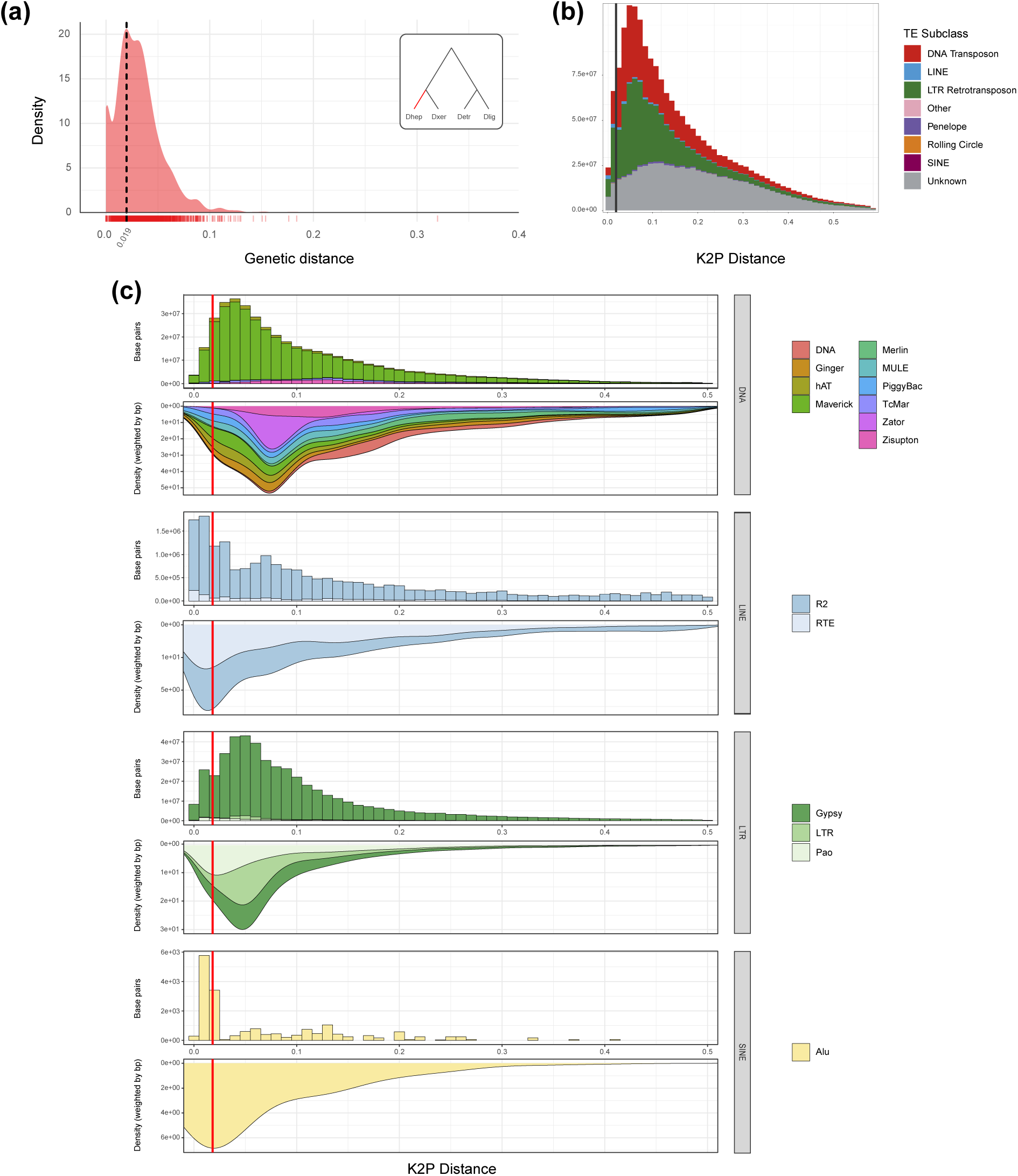
Density and repeat landscape analyses of *D. hepta.* **(a)** Density distribution of four-fold degenerate (4d) site genetic distances. The dotted vertical line indicates the distribution mode. **(b)** Repeat landscape plot summarizing relative TEs activity using Kimura 2-parameter (K2P) divergence; recent activity is shown on the left-hand side of the plot. The solid vertical line represents the 4d divergence mode for comparison. **(c)** Repeat landscapes of DNA, LINE, LTR, and SINE superfamilies. Upper plots represent cumulative base pairs grouped by divergence and family (color-coded), while inverted plots display the corresponding density distributions. Vertical red lines indicate the 4d divergence mode for reference.

### 2.4. Transposable element composition and analysis

We examined the repeat content across all pseudochromosomes (Figure 3a; Supplementary Fig. S6). Transposable elements (TEs) annotations revealed that a large portion of the repetitive content in *Dugesia* corresponds to DNA elements (50-65%), whereas in *S. mediterranea* this category accounts for less than 40% of the repeats (Supplementary Fig. S6a). In absolute terms, the increase in GS appears largely attributable to an expansion of TEs, particularly DNA/Maverick, DNA/DDE, and, as reported in *Schmidtea*, LTR/ Gypsy elements (Figure 3a; Supplementary Fig. S6). When comparing the relative composition of repeats, the proportion of DNA/DDE elements is similar between *Schmidtea* and *Dugesia*. However, *Dugesia* exhibits a clear expansion of DNA/Maverick elements, most notably in *D. etrusca*, which are the largest TEs known in eukaryotes, 15-40 kb^48,49^. Comparisons with the *S. mediterranea* genome also revealed that *Dugesia* and *Schmidtea* share a highly similar LTR composition, with > 90% of LTR TE being Gypsy elements, yet significant differences are found in the LINE and SINE orders (Supplementary Fig. S7d,e). In *Dugesia*, there is a clear expansion of LINE/RTE elements, the smallest non-LTR retrotransposons^50^, as well as an expansion of the SINE/5S and SINE/7SL subfamilies, which are seemingly absent in *Schmidtea* (Supplementary Fig. S6e). Notably, ∼65% of SINE in *D. hepta* are SINE/5S elements, 3∼4 times more than in *D. etrusca* or *D. liguriensis*.

We explored the pronounced expansion of SINE/5S elements in *D. hepta*, hypothesizing that it might indicate a role of TEs in CRs detected in the macrosynteny analysis. To test this, we examined the TEs density along *D. hepta*’s multisyntenic chromosome 2 using the R package *karyoploteR*^51^, and assessed whether any TEs family was overrepresented within 20kb windows flanking synteny breakpoints using the R package GenomicRanges v.1.46.1^52^. We found no particular TEs family to be significantly enriched or depleted near these breakpoints (Figure 3c,d; Supplementary Table S2).

We investigated the relative TEs activity period by estimating and comparing the distribution of the Kimura 2-parameter (K2P) divergence of TE subclasses and families of Dhep_v1’s assembly using EarlGrey^53^, against the distribution of four-fold degenerate (4d) sites divergence, which we used as a neutral model of *D. hepta*’s evolution, computed with phyloFit^54^. Our results show that the mode of most TE families and subclasses divergence was larger than that of the 4d divergence (Figure 5; Supplementary Table S3), suggesting that the relative activity of these TEs predates *D. hepta*’s speciation.

### 2.5. Gene family members’ gain, loss, and turnover rates

WGD events involve a rapid loss of gene copies. We explored this effect by estimating the gene gain and gene loss turnover rates using BadiRate v.1.35^55^. We first inferred the orthologous relationships with Broccoli between the four *Dugesia* species and *S. mediterranea*, yielding 18,719 orthogroups (OGs hereafter and referred to as gene families throughout the text). Moreover, we also included all singleton proteins, resulting in a total of 54,011 OGs, needed for the gene family analysis. We compared different branch-rate models under the birth, death, and innovation framework (BDI) of BadiRate. Our results supported a free-rates model (Supplementary Table S4) and showed that gene family gains were concentrated on the terminal branches of the phylogeny (Figure 3e), rather than on internal ones expected under a WGD scenario. This pattern is consistent with gene family turnover rates estimates, which increased in both birth and death rates along the external branches (Figure 3e; Supplementary Table S4), with the largest turnover rates present in the Corso-Sardinian clade.

## Discussion

### New genomes to contribute to the Tree of Life

Planarian genomes have long posed challenges for assembly owing to their extreme compositional bias, high repeat content, and mosaicism. This has limited the availability of high-quality genomic resources for Tricladida. The high-quality chromosome-scale assemblies presented here directly address this gap, providing a substantial advance.

These new *Dugesia* genomes are markedly more contiguous and complete than the reference available for the genus (cf. Tian et al.^32^), representing over a 200-fold improvement in continuity (Table 1). Although the sequence captured in pseudochromosomes is relatively high (Table 1; Supplementary Fig. S1, S2), overall completeness does not attain that achieved for *Schmidtea* (cf. Ivanković et al.^27^). However, these genomes represent a necessary advance toward improving the representation of neglected lophotrochozoan phyla in genomic databases, and constitute a first step toward developing a group-specific BUSCO set for Platyhelminthes in the future. Our results, nevertheless, should be contextualized in light of the well-known genomic features of planarians: extreme compositional AT-bias (>70% A/T), high repetitive content, and the characteristic mosaicism in *Dugesia* (see mosaic-Meselson effect^33^). These features likely complicate the haplotig collapsing and contribute to the inflated numbers of scaffolds. Indeed, all *Dugesia* assemblies shared a similar high fraction of repeat content (Table 2) that substantially exceeds previously reported values in *D. japonica* (∼65% ^32^), *S. mediterranea* (∼62% ^42^), or *Schmidtea polychroa* (Schmidt, 1861) (∼58% ^56^). All these features reinforce the findings of Ivanković et al.^27^, highlighting that planarian genomes remain technically challenging to assemble. In fact, in *D. etrusca* and *D. xeropotamica,* the proportion of the sequence captured in pseudochromosomes was lower than in the other two *Dugesia* species. However, when these proportions are evaluated against their flow cytometry-based GS estimates (Table 1), they increase markedly. This further underscores the technical challenges inherent to assembling planarian genomes using current methodologies.

### CRs in *Dugesia* and the origin of *D. hepta*’s karyotype

The availability of these new *Dugesia* assemblies provided, for the first time, the opportunity to explore macrosynteny relationships between genera as well as to assess the origin of the peculiar haploid complement of *D. hepta* (n = 7), a notable exception to the predominant regional set of n = 8^57^ (Figure 1). Our results showed several translocations involving chromosomes 5, 6, 7, and 8 between the Corso-Sardinian and the Iberia-Apennines-Alps clades, whereas the four largest chromosomes displayed a well-conserved synteny (Figure 3b). CRs had also shaped *D. hepta*’s karyotype, where we detected relevant translocations and pericentric inversions, particularly in chromosomes 2, 3, and 7.

Chromosome 2 is of special interest, as it is key to explaining the reduction in haploid chromosome number. CRs have long been recognized as a major force in genome evolution, with a relevant role in speciation^58,59^. Specifically, changes in chromosome number can often lead to reproductive isolation due to heterokaryotypic individuals experiencing reduced fertility (or overall reduced fitness) caused by meiotic segregation problems. In this context, the case of *D. hepta* is unambiguous. Previous genomic analyses showed no evidence of past or present gene flow between *D. hepta* and its congenerics^37^, co-occurring in the same localities. While it is difficult to determine whether *D. hepta*’s CRs were concomitant with or followed its speciation, they likely played a subsequent role in reinforcing its reproductive isolation. It is well established that TEs can contribute to the origin of CRs (reviewed in ^60^). Evidence for the role of CRs in the speciation could come from linking TEs insertions to CRs breakpoints and the demonstration of a concomitant activity of these TEs with the speciation event. However, our results did not identify any TE family significantly enriched in chromosome 2’s synteny breakpoints. Furthermore, our estimates of the relative TEs activity (Figure 5) suggested that most TEs families’ mobilization likely predated *D. hepta*’s speciation. Although LINE/R2 and SINE/Alu elements show activity periods concomitant with the speciation event, this evidence is insufficient to implicate them as drivers of the CRs that gave rise to chromosome 2, given the small number of base pairs involved and the absence of enrichment signals (Figure 3d; Figure 5c).

### Genome size evolution in planarian flatworms

Little is known about the evolutionary dynamics of GSs in planarian flatworms. A recently published study by Ivanković et al.^27^ provided the first comparative view of four species within the genus *Schmidtea*, revealing substantial differences between a clade of small GSs (*S. mediterranea* and *S. polychroa*; 0.82∼0.78 Gb, respectively) and a clade of relatively large GSs (*Schmidtea lugubris* Schmidt, 1861 and *Schmidtea nova* Benazzi, 1982; 1.5∼1.2 Gb, respectively). The authors proposed that a large portion of this increase could be attributed to TE expansions and an increased length of the protein-coding genes caused by TE insertions within introns.

Incorporating our *Dugesia* results into the comparative framework provides new insights into planarian GS evolution. Many Western Palearctic *Dugesia* species share a haploid complement of n = 8, and GS estimates for Western Mediterranean species indicate that ∼2 Gb is a common size (e.g., *D. benazzii*, 2.05 Gb; *D. leporii*, 2.38 Gb). Owing to the lack of GS estimates for other dugesiid genera, including *Girardia* Ball, 1974, or *Recurva*, it remains difficult to ascertain whether the observed differences reflect a genome expansion in *Dugesia* or a genome reduction in *Schmidtea*. Nevertheless, when comparing the available Dugesiidae GS estimates in the Animal Genome Size Database^61^ (http://www.genomesize.com, last accessed in October 2025; cf. *Dugesia subtentaculata* (Draparnaud, 1801), 1.66 Gb; *Girardia tigrina* (Girard, 1850), 1.40∼1.88 Gb) with our estimates (Table 1) and those reported previously^27^, the data are more consistent with a lineage-specific genome expansion in Western Mediterranean *Dugesia*, potentially coupled with a genome reduction in the *mediterranea-polychroa* clade.

Bearing in mind the aforementioned differences between *Dugesia* and *Schmidtea,* we contemplated the hypothesis of a WGD event as the underlying cause not only for the karyotypic differences but also for the increase of GS in *Dugesia*, in contrast to the proposed TE expansion-driven way of increase within *Schmidtea*. Yet, our results consistently refute the WGD hypothesis. The distribution of *K*_s_ values showed exponential decay (Figure 2a), as expected under the assumption of constant gene gain and loss events, without any WGD hallmark^62^. Likewise, collinearity, macrosynteny, and gene family members’ gain/loss analyses also support this conclusion. WGDs are expected to leave large blocks of intragenomic collinearity^63^, and in *Dugesia*, collinearity is negligible in our genomes. For instance, autopolyploids such as *Arabidopsis thaliana* can retain up to ∼27% collinear genes^45^. Furthermore, we observed neither an increase in turnover rates at the base of the genus *Dugesia* (Supplementary Table S4) nor multimatching syntenic blocks in *Schmidtea* (Figure 3b). Together, these observations indicate that the larger GS in *Dugesia* arose from mechanisms other than WGD.

The analysis of TEs revealed that the expansion of multiple TE families in *Dugesia*, as previously reported in the *lugubris-nova* lineage^27^, likely underlies the observed increase in GS. However, the specific families involved differ, with DNA/Maverick and DNA/DDE families showing the most pronounced expansions. In contrast to *Schmidtea*, we did not detect an increase in intronic lengths attributable to TEs insertions. Together, these observations indicate that TE expansions have contributed to genome increases in *Dugesia*, but that the specific TE families involved, and potentially the evolutionary forces driving their proliferation, are not shared across species.

The evolutionary context of the Corso-Sardinian clade is conditioned by the insularity of its populations, and the largest flow cytometry GS estimates in *Dugesia* come from this group. Two of the main mechanisms that may contribute to GS increase are (i) changes in ploidy (ii) and TEs mobilization^60,64^. According to the mutational hazard hypothesis (see Lynch et al.^65^), the fate of DNA gains, including TEs expansions, is governed by the mutation rate and the effective population size (*N*_e_). Indeed, in insular systems, smaller *N*_e_ are expected, leading to relaxed selective constraints genome-wide^66,67^ (i.e., reduced efficacy of selection and predominance of genetic drift). Under this scenario, TE dynamics may shift toward accumulation, with slightly deleterious TE insertions persisting because purifying selection cannot efficiently purge them. *Dugesia hepta* and *D. xeropotamica* are both endemics from Sardinia and Corsica with very restricted ranges of distribution, and coincidentally, both exhibit the largest genomes according to flow cytometry estimates (Table 1). Moreover, the largest GS estimate known for *Dugesia* belongs to a Sardinian endemic as well, *D. leporii* (2.38 Gb), further reinforcing this pattern. The evolutionary histories of these species are tightly linked to some of the major paleoenvironmental shifts in the Western Mediterranean^37^, and it is reasonable to assume that both species experienced fluctuations in *N*_e_, including likely bottlenecks occurring during the Pleistocene glacial phases on both islands^68,69^. Consistent with this view, comparisons of 4d distances of *Dugesia* did not show significant differences among *Dugesia* species or clades (Supplementary Fig. S8), indicating that the neutral evolutionary rates are broadly similar. This, in turn, strengthens the idea that differences in GS might instead reflect historical variations in *N*_e_. Such demographic fluctuations in the insular lineages should have an impact on gene family turnover rates, and indeed, our analyses reveal a 1.3∼2.5-fold and a 2.6∼3.8-fold increase in gene birth and death turnover rates, respectively, in *D. hepta* and *D. xeropotamica* relative to the continental counterparts (Supplementary Table S4).

### Planarian genome architecture: resilience in front of massive genome scrambling?

A further striking finding is the discrepancy in structural genome stability observed between *Dugesia* and the patterns previously reported for *Schmidtea* (cf. Ivanković et al.^27^). Our results revealed that structural genome rearrangements are widespread between the two genera, resulting in a pattern of near-complete genome scrambling (Figure 3b). In *Dugesia*, macrosynteny analysis showed that two major fusion events underlie the haploid chromosome complement of *D. hepta* (n = 7), as well as several translocations affecting large chromosomal fragments between the two major clades, a phenomenon previously reported in other *Dugesia* species^70,71,72,73^, and also frequent in fissiparous individuals^74^. Barring that, the four studied *Dugesia* species retain a largely conserved macrosynteny, despite their deep divergence time (∼30 Mya^26^; Figure 1), in stark contrast to the extensive syntenic disruption observed in *Schmidtea*. Based on the patterns reported in the latter genus, it has been proposed that pervasive structural genome instability is a general feature of flatworm genome evolution^27^. However, our findings in *Dugesia* suggest that this feature may not be generalizable. Nevertheless, divergence-time estimates for *Schmidtea* species are currently lacking, leaving open the question of whether their extensive CRs represent a genus-specific feature, a broader characteristic of Dugesiidae, or instead a signature of triclad flatworm lineages that diverged a long time ago. Assuming comparable substitution rates, the lower divergence at four-fold degenerate sites in *Dugesia* (0.06 substitutions per site from their MRCA) relative to *Schmidtea* (∼0.3, cf. Ivanković et al.^27^) suggests that the MRCA of the studied *Schmidtea* species predates that of the Dugesia included in this study. It would be most interesting to obtain chromosome-scale genomes of Malagasy *Dugesia*, the earliest diverging clade within the genus (>100 Mya^26^), to assess whether synteny is preserved only at ‘shallow’ evolutionary timescales in planarian flatworms. Nevertheless, our observation of a complete absence of synteny with the MALGs in *Dugesia* (Figure 4) does not contradict the hypothesis of the pervasive structural instability of the flatworm genome evolution proposed by Ivanković et al.^27^.

Taken together, our findings, along with those of Ivanković et al.^27^, raise questions about the causes and consequences of such extreme genome scrambling in triclad flatworms. While pervasive genome scrambling could potentially disrupt gene regulation and be detrimental, the observation that *S. mediterranea* exhibits tight, proximity-based associations between regulatory elements and their target genes points to a mechanism that may mitigate the negative effects of pervasive CRs. This mechanism may also explain why CRs are so frequent in the wild in fissiparous populations^74^. A similar link between compact regulatory structure and extensive genomic rearrangements has been proposed in tunicates and clitellates^21,75^. But how to explain this pronounced tendency to undergo chromosomal rearrangements? This genomic architecture could actually be a consequence of a constraint linked to one of the most relevant characteristics of planarians: their remarkable regenerative capacity and cell turnover. Many of their striking evolutionary features, such as the mosaic-Meselson effect^33^ and the persistence of ancient asexual lineages^35,36^, are intrinsically tied to the accumulation of mutational load and the hurdles of Muller’s ratchet^76,77,78^, and somehow planarians are able to circumvent these limitations^36,74^. High GC content has been shown to prevent double-strand breaks in prokaryotes^79^, yet it is also positively correlated with higher mutation rates^80^. Planarians, which exhibit exceptionally low GC content^27,56,81^ (Table 1), may have evolved this genomic feature as a compensatory strategy to reduce the burden of non-synonymous mutations accrued through their continuous regeneration and cell turnover. This could result in a genome composition that is resilient to mutational load, yet CRs-prone. Remarkably, *Hydra* (phylum Cnidaria), which exhibits regenerative abilities comparable to those of freshwater planarians, also shows an extreme compositional AT-bias^82^. Under this hypothesis, selective pressures may have favoured the emergence of a gene-centric regulatory architecture capable of mitigating the negative effects of such pervasive structural instability.

## Materials and methods

### Samples

All animals used for the analyses were collected from wild populations and reared in laboratory in 1× Montjuïc water (1.6 mM NaCl, 1.0 mM CaCl_2_, 1.0 mM MgSO_4_, 0.1 mM MgCl_2_, 0.1 mM KCl, and 1.2 mM NaHCO3 in DI H_2_O, pH 6.9–8.1) at a constant temperature of 14 °C. *Dugesia benazzii* samples were collected at 40.57498, 9.6746543, near Siniscola, Sardinia (Italy). *Dugesia etrusca* samples were collected at 43.497778, 10.625833, near the town of Rivalto (Italy). *Dugesia hepta* samples were collected at 40.756564, 8.604074, near the city of Sassari, Sardinia (Italy). *Dugesia leporii* samples were collected at 39.436662, 8.5082993, near the town of Fluminimaggiore, Sardinia (Italy). *Dugesia liguriensis* samples were collected at 43.823611, 6.581944, near the village of La Garde (France). *Dugesia xeropotamica* samples were collected at 42.473832, 8.795403, at the Lioli rivulet. Animals were starved for a week prior to DNA/RNA extractions. Animals destined for DNA extractions were flash-frozen in liquid nitrogen and stored at -80 °C before being sent for sequencing. Final species selected for genome sequencing were chosen based on overall extraction yield and quality.

### Flow cytometry

GS estimation was carried out by means of flow cytometry. Samples were prepared by incubating living individuals in a solution of 2% N-acetyl-L-cysteine at pH 7 to remove the mucus and minimize the formation of cell aggregates. Animals were then rinsed for 1 minute using a mixture of distilled water and tap water (1:1) and placed in separated Petri dishes. Thereafter, samples were minced using a sterilized stainless steel razor and incubated in 1 ml Galbraith’s isolation buffer^83^ supplemented with 10 μl of RNase A (1 mg/μl) for 15 minutes. After incubation, the cells were retrieved and separated by gently pipetting the liquid contents through a nylon mesh (pore size, 75 μm) into an Eppendorf tube. Further, the cell suspension was stained for 15 minutes with 30 μl of propidium iodide (stock, 1 mg/ml). The quantity of DNA was measured with a Gallios Flow Cytometer at the Unitat de Citometria dels Centres Científics i Tecnològics de la UB (CCiT, UB). Finally, GS was determined by comparing the readings of fluorescence of our samples against those of males of *Blatella germanica*, with a known GS of 2.025 Gb.

### DNA/RNA extractions, sequencing, and genome assembly

RNA extractions were performed using Trizol (Thermo Fisher Scientific, USA) following the manufacturer’s instructions. Total RNA quantification and integrity were assessed with a Qubit and a Bioanalyzer in the Centres Científics i Tecnològics, Universitat de Barcelona (CCiT, UB). Truseq stranded and ribo-zero libraries were constructed in Macrogen Inc., (Macrogen Europe, Madrid) to obtain Illumina paired-end sequencing.

DNA extraction, sequencing, and genome assembly were performed at Dovetail Genomics (Scotts Valley, CA). High-quality genomic DNA of each species was extracted from pooled samples following the protocol described in Zhang and Alvarado^84^. Genomic sequence data were generated using a single library per species: PacBio CLR (*D. hepta, D. etrusca*) and PacBio HiFi (*D. liguriensis, D. xeropotamica*) libraries. Omni-C libraries (one library per each species) were then built and sequenced using Illumina 150-bp paired-end (HiSeq X platform) to perform the chromosome-level scaffolding using the HiRise software^85^. The assemblies’ completeness was assessed by applying the pipeline of Benchmarking Universal Single Copy Orthologues (BUSCO; v.5.4.2^41^), searching the metazoa (odb10; 954 genes), arthropoda (odb10; 1013 genes), and eukaryota (odb10; 255 genes) data sets.

### Contaminants sequence removal

Mitochondrial genome sequences were removed from the assembled scaffolds using BLAST+ blastn v.2.12.0^86,87^ searches against the mitogenomes of *D. japonica* (GenBank accession number AB618487) and *Dugesia. ryukyuensis* Kawakatsu, 1976 (AB618487). Any hit of >1kb was removed and replaced by Ns in the corresponding scaffold. We then filtered out potential contaminant sequences and assembly artifacts using several criteria. First, we mapped each assembly against the NCBI nucleotide database using BLAST+ blastn and Blobtools v.3.4.0, as implemented in BlobToolKit^88^, and discarded scaffolds with bacterial, virus, or fungal hits. Next, we calculated the mean depth of coverage for the remaining scaffolds and discarded those falling outside the 95% confidence interval that also lacked BUSCO hits.

### Identification of repetitive elements (RE)

We analysed repetitive regions of the genome assemblies using a combination of *de novo* TEs identification with RepeatModeler v2.0.3^89^ and DeepTE^90^, and similarity-based searches using RepearMasker v.4.1.2^91^. We built a RE database for each species. We modelled RE using RepeatModeler and parsed its output to classify RE into identified and unknown elements. Thereafter, we used DeepTE to classify the unknown RE using the metazoan model (-sp M). Lastly, all classified RE were concatenated into a single database to be used to soft-mask each genome assembly with the `–xsmall` option of RepeatMasker. Additionally, we performed *de novo* repeat identification in *D. hepta* using EarlGrey^53^ to estimate the repeat landscapes of the major TE families to assess their relative activity in *Dugesia* genomes.

### Structural and functional genome annotation

Structural annotation of the soft-masked genome assemblies was performed with BRAKER2^92,93,94,95^, using species-specific RNAseq data that had been previously mapped to each assembly using STAR v.2.7.10b^96^. Functional annotation of the resulting gene models was carried out using BLASTP v.2.2 (E-value = 10^-3^) searches against Swiss-Prot, complemented with searches against the eggNOG database, using eggNOG-mapper v.2.1.9^97,98,99^, and protein domain detection with InterProScan v.5.56.89^100^. All functional evidence was integrated. Concurrently, transfer RNA genes (tRNAs) were identified using the tRNAscan-SE v2.0.7^101^.

### Intron and CDS lengths

We estimated intron and CDS length distribution of *Dugesia* and *S. mediterranea* using only single copy orthologs (SOs; 1:1:1:1:1) from the inferred orthologous groups (OG) determined with Broccoli v.1.2^46^. For the analysis, we used information from the largest scaffolds or pseudochromosomes (*D. etrusca*, *D. liguriensis*, and *D. xeropotamica*, 8 scaffolds; *D. hepta*, 7 scaffolds; *S. mediterranea*, 4 scaffolds, Guo et al. (2022) assembly). Intron and CDS length distributions were visualised using *ggplot2*.

### 4-fold degenerate sites analysis

We estimated the putative neutral divergence using four-fold degenerate (4d) sites. For that, first, we extracted these sites from the coding sequences (CDS) of single-copy *Dugesia* orthologues identified by Broccoli, in combination with BRAKER’s structural annotations. Each orthogroup was aligned with TranslatorX^102^, using its implementation in MAFFT^103^ to maintain the correct reading frame and ensure positional homology across nucleotides. Finally, we used phyloFit^54^ with the default REV substitution model to estimate the neutral model from the 4d sites across all alignments, and visualized the resulting distribution of genetic distances using *ggplot2*.

### Detection of whole-genome duplication

We constructed the *K*s distribution to evaluate whether a WGD event could underlie the conspicuous GS differences between *Dugesia* and *S. mediterranea*. First, using wgd v2^44^, we inferred the paralogous gene families for each *Dugesia* species from their coding sequences (`wgd dmd`). Thereafter, we estimated (`wgd ksd`) and visualized (`wgd viz`) the *K*s distributions. Furthermore, we identified all potential homologous protein pairs within each species using BLASTP v.2.2 (E-value = 10^-10^ and top 5 matches), and used MCScanX^45^ to detect collinear gene pairs.

### Macrosynteny analysis

To explore potential CRs in *D. hepta*, we used the R package GENESPACE v.1.2.3^47^, which combines OrthoFinder^104^ and a reimplementation of the MCScanX algorithm to infer gene-ordered-based syntenic blocks, and visualized them using the built-in riparian plot function. Additionally, we applied the ODP tool^105^ to determine whether ancestral metazoan linkage groups (MALGs), highly conserved linkage groups across bilaterians, cnidarians, and sponges^11^, are preserved in *Dugesia* genomes. This tool uses hidden Markov models to identify homologs of MALG proteins and applies Fisher’s exact test to determine their significant enrichment on specific chromosomes.

### TEs enrichment analysis at syntenic breakpoints

We visualized the density distribution of TE along chromosome 2 of *D. hepta* using the R package *karyoploteR*^51^ to identify regions potentially involved in CRs detected in the macrosynteny analysis. In addition, we tested whether specific TE families were over-represented nearby synteny breakpoints on the same chromosome using the R package *GenomicRanges* v.1.46.1^52^. Based on GENESPACE syntenic blocks delineation and RepeatMasker’s annotation, we established flanking windows of 20kb surrounding each breakpoint and counted all TEs present within these regions. To assess enrichment, we generated a null distribution of repetitive sequences by placing an equal number of randomly distributed windows along the chromosome and then computed 1000 permutation iterations. A two-tailed test was performed to assess whether TE families were significantly enriched or depleted in the flanking windows relative to the null distribution. To account for multiple testing and low-abundance families, we applied two filtering steps: (i) p-values were calculated only for TE families with an average expected count of 10 or more sequences, and (ii) the resulting raw p-values were adjusted using the Benjamini-Hochberg false discovery rate method.

### Estimation of gene gain and loss

We estimated gene gains and losses using BadiRate v.1.35^55^, under the birth, death, and innovation model (BDI), applying a maximum likelihood (ML) method to estimate the turnover rates. Furthermore, we also tested multiple branch models to estimate the best-fitting one by comparing the Akaike Information Criterion (AIC) values. The models considered were: (i) global rates (GR), in which all branches evolve under the same turnover rates; (ii) free rates (FR), which allows each branch to evolve under distinct rates; and (iii) branch-specific (BR), where a subset of branches share different turnover rates from the background (here, those leading to *Dugesia*). We used the OG counts obtained from Broccoli and a time-calibrated tree based on the SOs set. Given that Broccoli may leave some proteins unassigned to OGs when handling complex evolutionary scenarios, we also included the singleton OGs for this analysis (eg., 1:0:0:0:0). Since BadiRate is prone to crash in the analysis of highly heterogenous counts, we analyzed these cases (only 0.02% of the OGs) separately using the best-fitting branching model using Wagner parsimony. For the analysis, we inferred a time-calibrated tree with IQ-TREE v2.4.0^106^ using an admixture model (LG+C60+F+G+R3) and subsequently calibrated with the *chronos* function from the R package *ape*^107^. We used four calibration points: (i) the *Schmidtea-Dugesia* split (130-160 Mya), (ii) the most recent common ancestor (MRCA) of the featured *Dugesia* (23-36.5 Mya), (iii) the MRCA of the *hepta-xeropotamica* (4.25-8.87 Mya) and (iv) the MRCA of *etrusca-liguriensis* (6.3-13.3 Mya)26,37.

## Acknowledgements

We are thankful to Longhua Guo and Xinghua Li for providing some of the necessary data for *Schmidtea mediterranea* to carry out some of the analyses of this study. We also express our sincere gratitude to Gema Blasco for her dedication and exceptional care when rearing the planarians in the laboratory. We thank Alejandro Sánchez-Gracia and Sara Guirao-Rico for their invaluable inputs throughout the course of the study’s elaboration.

## Author contributions

J.R., and M.R. designed the study. D.D.S., I.T.A., and M.R. carried out sampling efforts. D.D.S. and I.T.A. performed the DNA extractions and the main comparative genome analyses. V.A.P. and M.O.M. provided bioinformatics assistance and helped perform some of the analyses. D.D.S. wrote the first version of the manuscript and conceived all data visualization. J.R. and M.R. supervised the research and final draft of the manuscript.

## Supplementary Information: Figures

**Supplementary Fig. S1.**
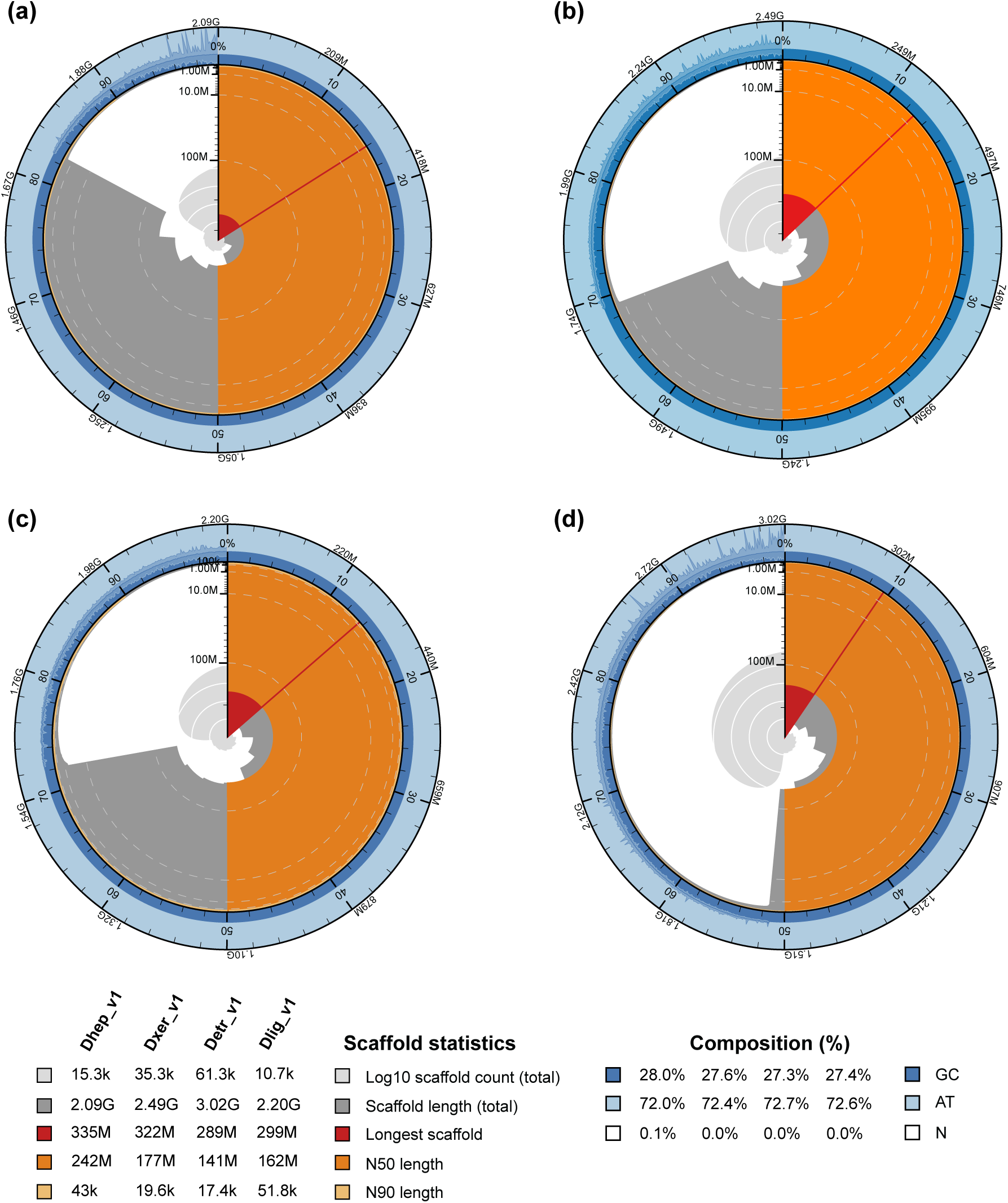
Snail-plots summarizing genome assembly statistics for (a) *D. hepta*, (b) *D. xeropotamica*, (c) *D. liguriensis*, and (d) *D. etrusca*.

**Supplementary Fig. S2.**
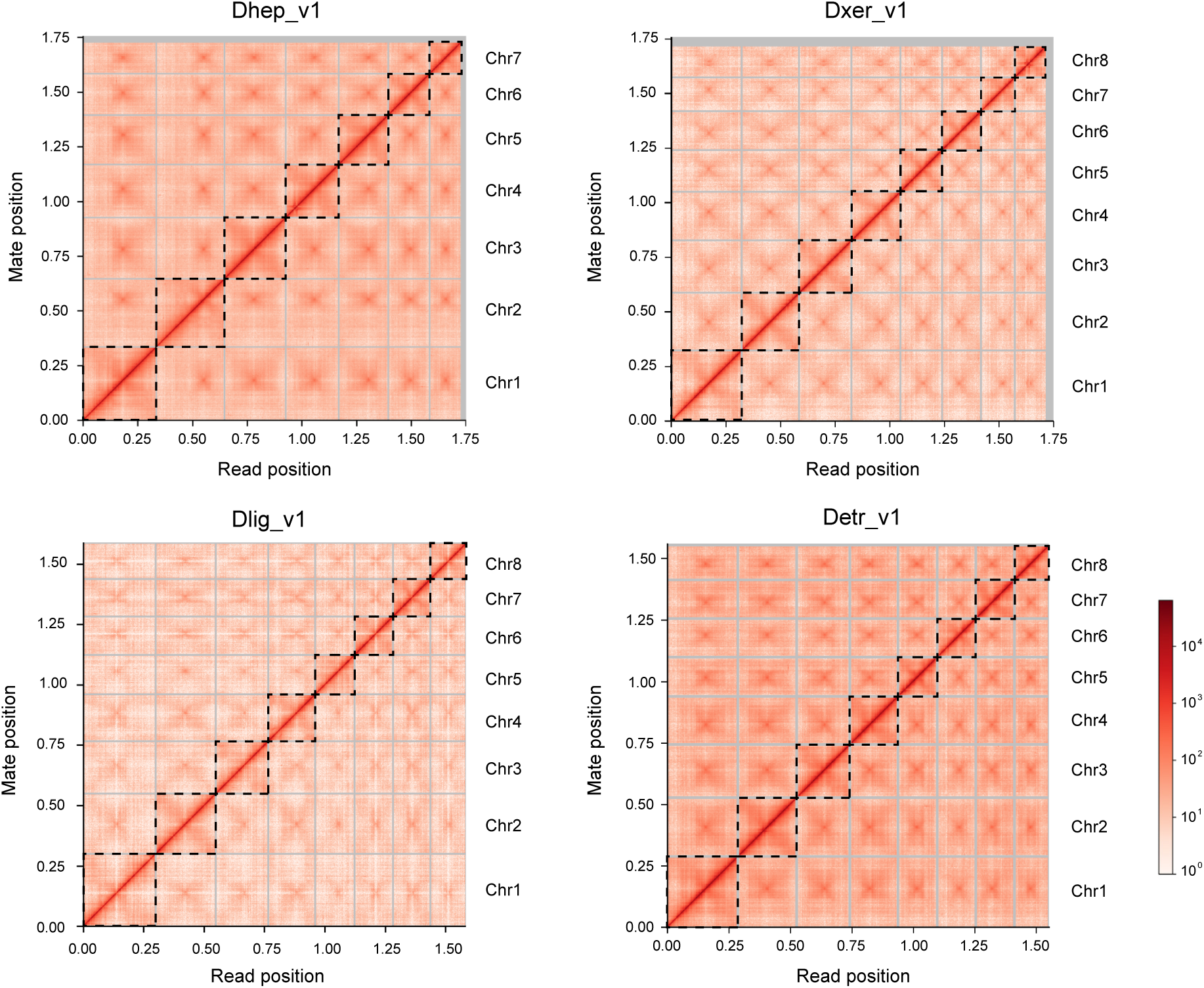
Long-range contact heatmap of paired Omni-C reads depicting the major scaffolds of

**Supplementary Fig. S3.**
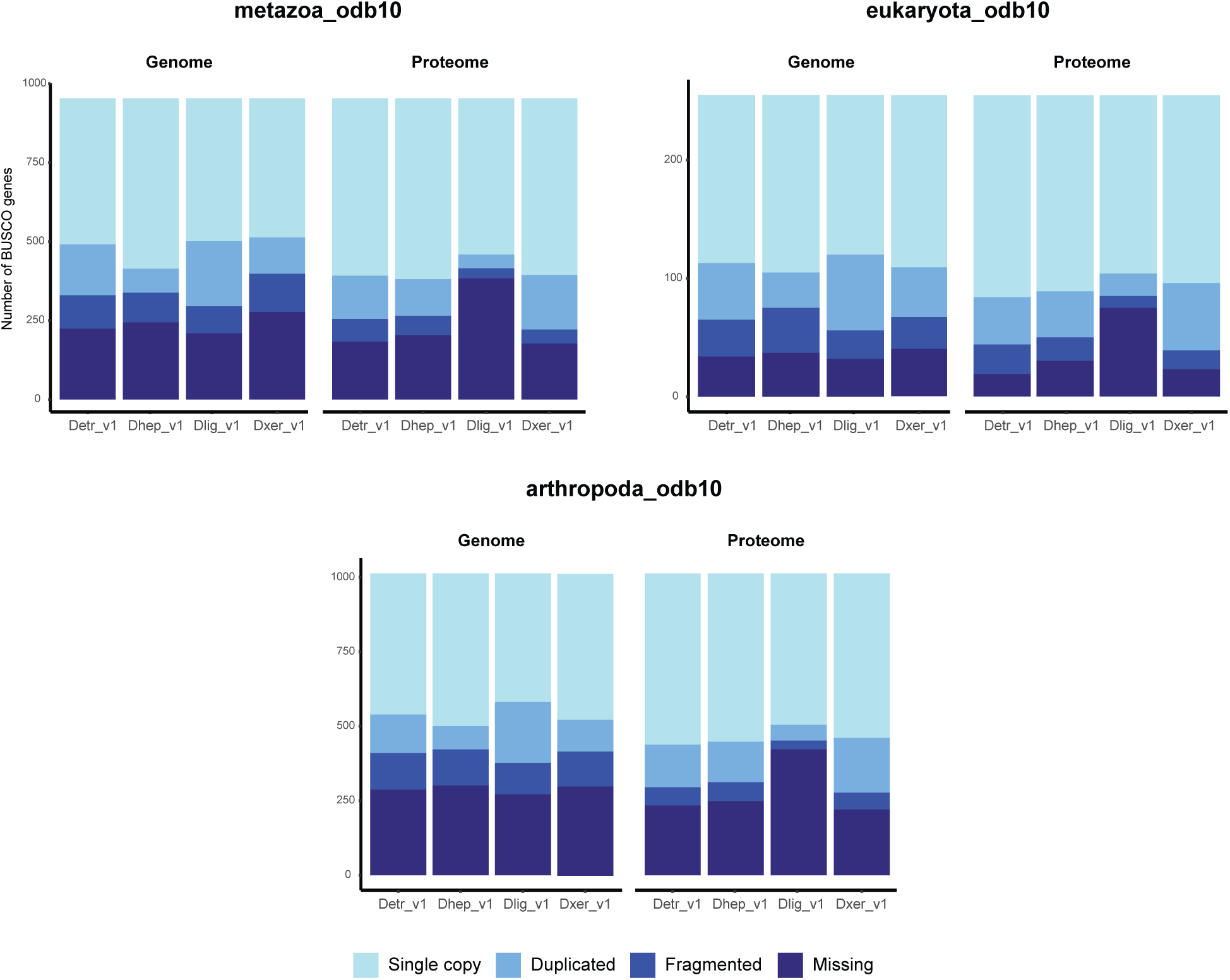
BUSCO assessment of the genome assemblies before (Genome) and after (Proteome) functional annotations.

**Supplementary Fig. S4.**
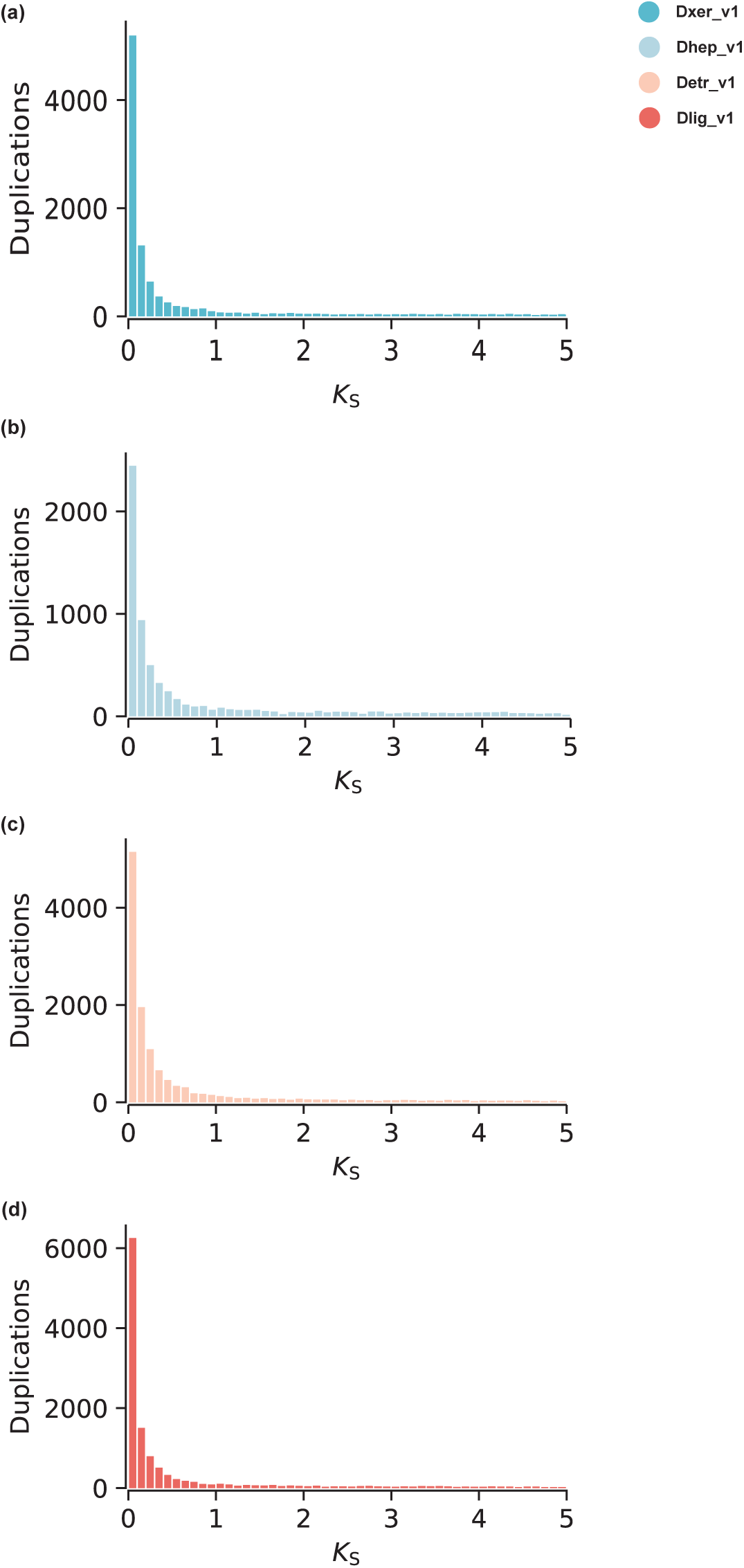
Paranome *K_s_* distributions of the *Dugesia* genome assemblies showing a pattern of exponential decay.

**Supplementary Fig. S5.**
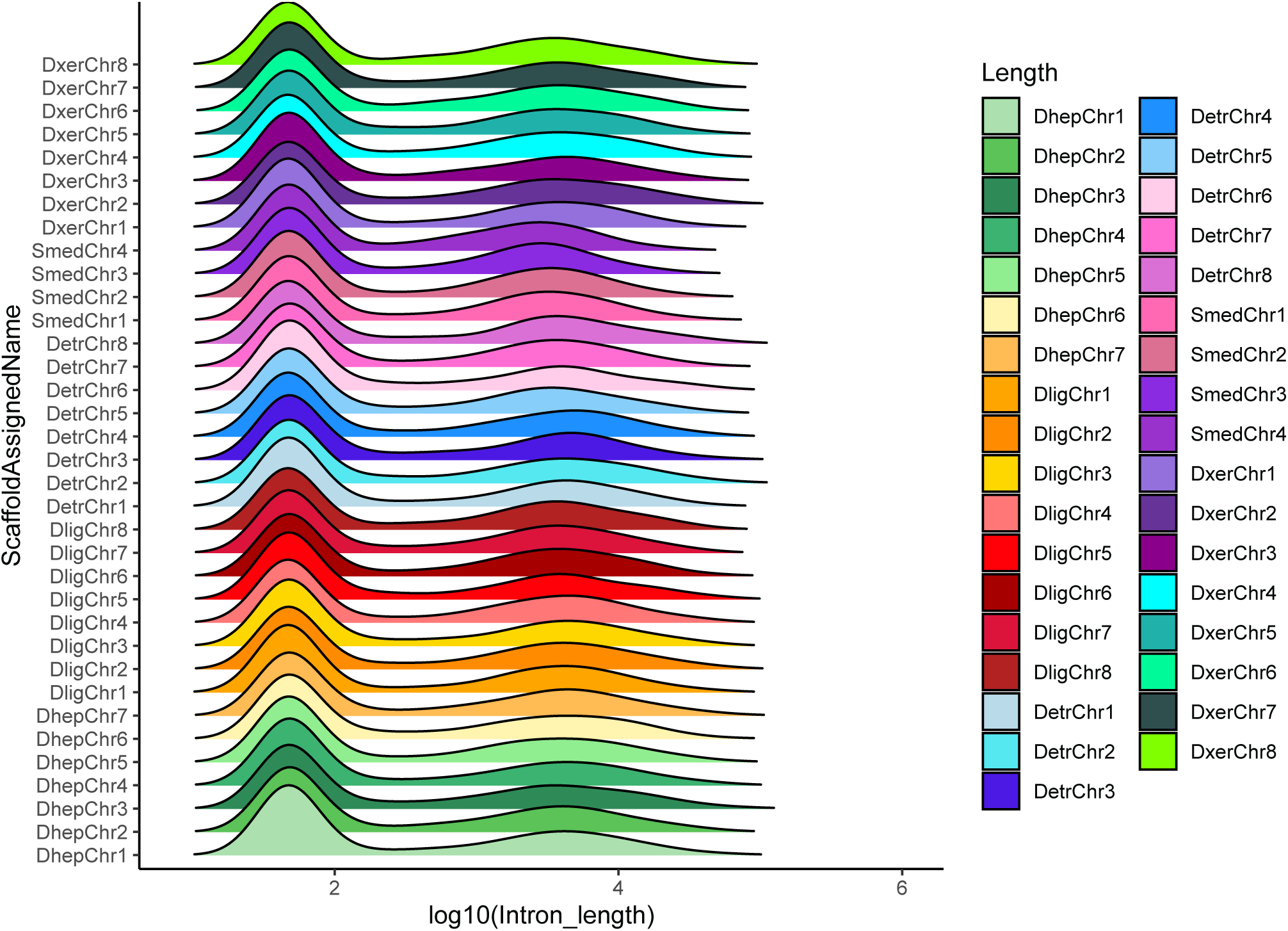
Ridgeline plots showing the density distribution of intron lengths of all introns comprised in the 2266 single-copy orthogroups shared between *S. mediterranea* (Smed), *D. hepta* (Dhep), *D. xeropotamica* (Dxer), *D. etrusca* (Detr) and *D. liguriensis* (Dlig) separated by chromosomes.

**Supplementary Fig. S6.**
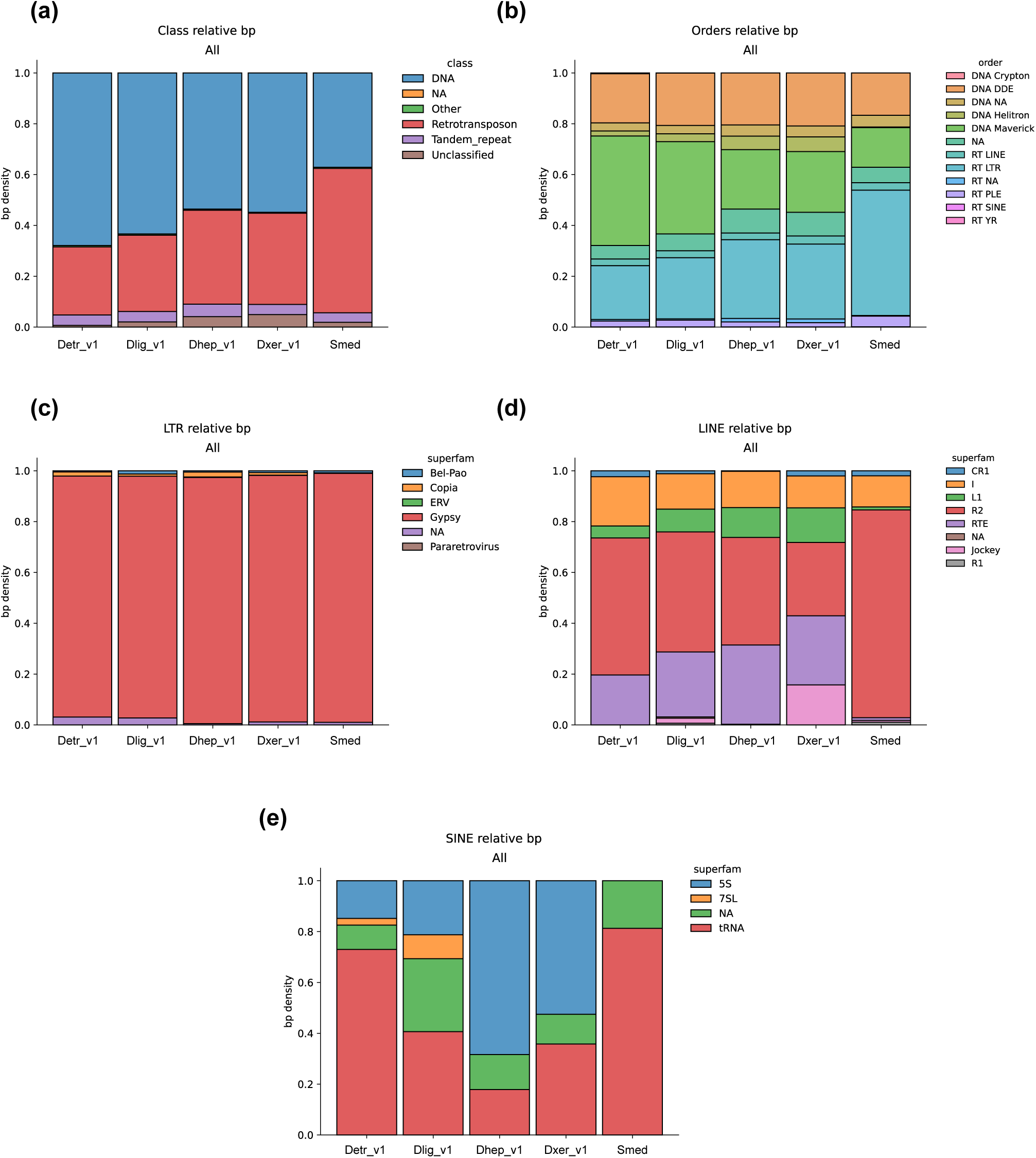
Comparison of the relative contributions (%) of each class (a) and order (b) of repetitive elements to the repeat content of each genome assembly. (c) Relative superfamilies contributions of repetitive elements to the LTR retrotransposon elements. Relative superfamilies contributions of repetitive elements to the (d) LINE retrotransposon and (e) SINE retrotransposon elements.

**Supplementary Fig. S7.**
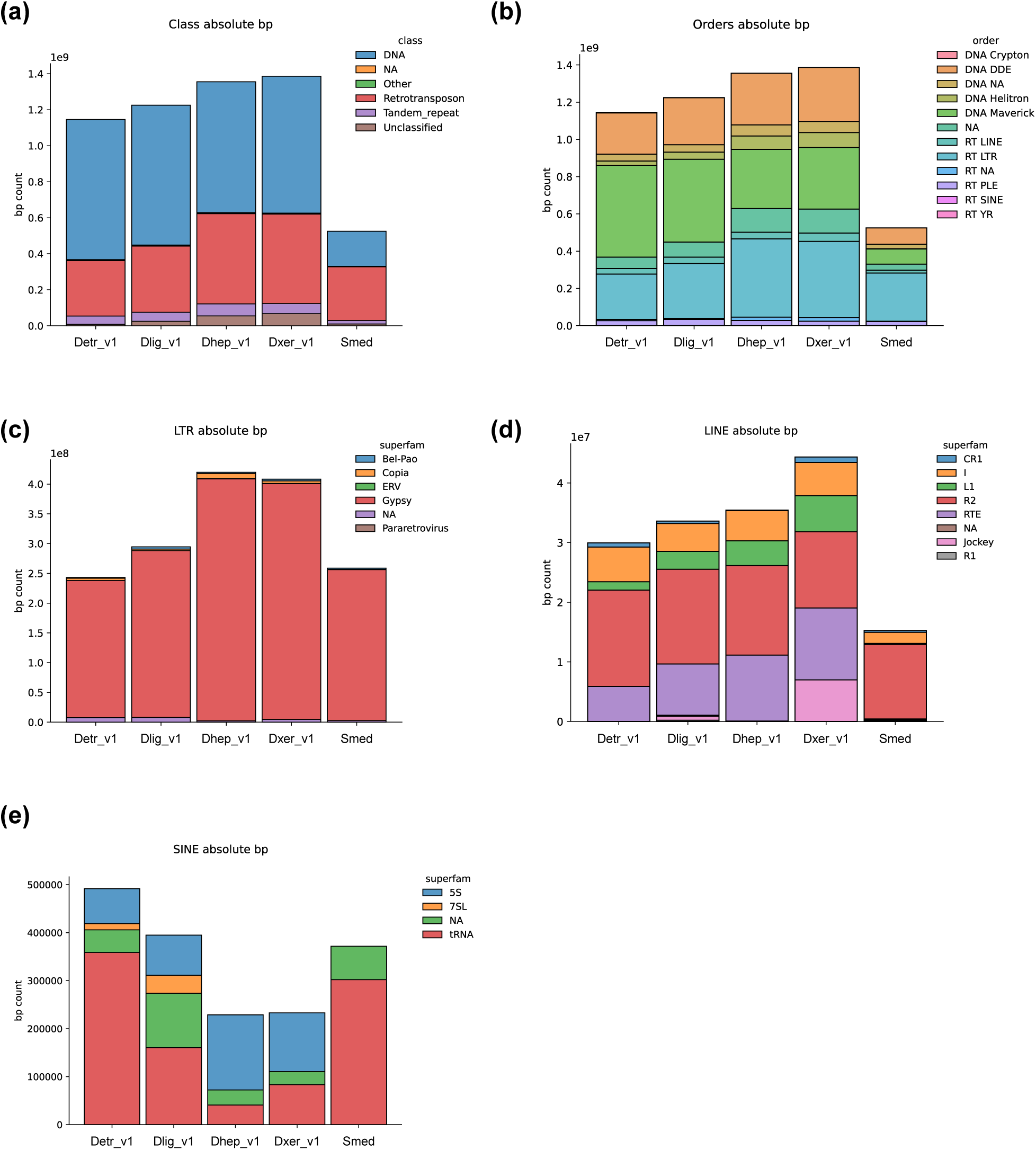
Stacked histograms showing the total amount (bp) of repeat content of each genome assembly classified into (a) classes of repetitive elements and (b) orders within said classes. (c) Close up look at the proportion of LTR retrotransposon elements. (d) Close up looks at the proportion of LINE retrotransposon elements and (e) SINE retrotransposon elements.

**Supplementary Fig. S8.**
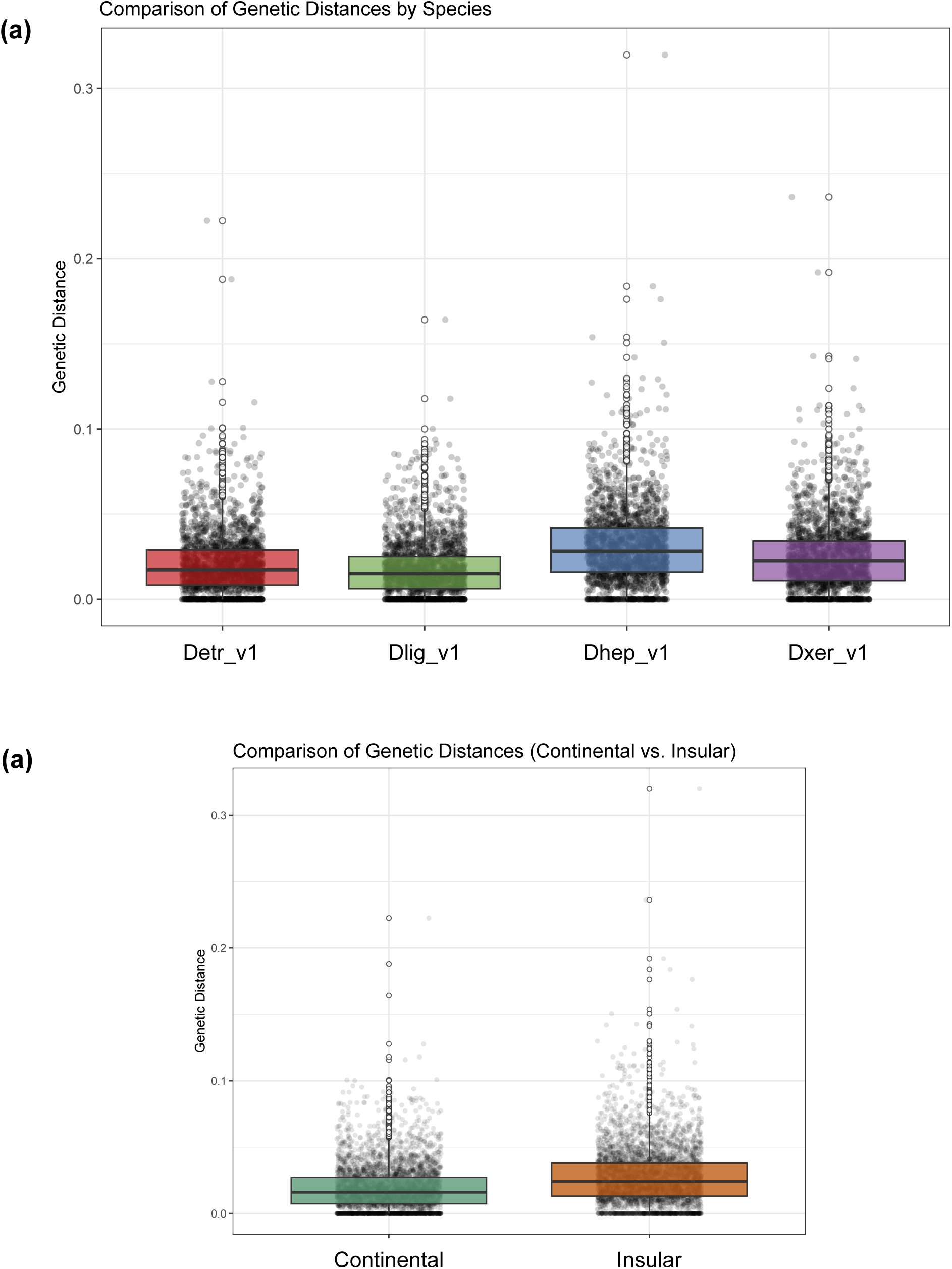
Comparison of the distributions of four-fold degenerate (4d) distances per species (a) and grouping continental and insular lineages (b).

